# Mechano-sensing role of oncogenic lncRNA NEAT1 on soft vs. stiff substrates and its implications for glioblastoma progression

**DOI:** 10.1101/2024.11.20.624463

**Authors:** Arpita Ghosh, R. Soundharya, Mohit Kumar Jolly, Abhijit Majumder

**Affiliations:** Department of Chemical Engineering, IIT Bombay, Powai, Mumbai 400076, India; Department of Bioengineering, Indian Institute of Science, Bengaluru 560012, India

## Abstract

Long non-coding RNAs (lncRNAs) have gained increasing recognition as key regulators of cancer biology, overturning their earlier classification as non-functional genomic elements. Among them, NEAT1 (Nuclear-Enriched Abundant Transcript 1) has emerged as a prominent oncogenic lncRNA involved in multiple solid tumors, including liver, prostate, and gastric cancers, as well as gliomas. Although the role of NEAT1 in cancer has been extensively studied, most of these investigations have focused on biochemical factors, leaving the mechano-sensitivity of NEAT1 to mechanical alterations in the tumor microenvironment (TME) largely unexplored. As tumors progress, the TME undergoes significant mechanical changes, particularly extracellular matrix (ECM) stiffening, which modulates cellular behavior through mechano-transduction pathways. While much research has focused on how these mechanical cues influence protein-coding gene expression, their impact on lncRNAs such as NEAT1 remains understudied. This study aims to address this gap by investigating the mechano-sensitivity of NEAT1 in glioblastoma (GBM) cells cultured on polyacrylamide (PAA) gels that mimic the brain’s physiological stiffness (0.5 kPa) versus conventional, stiff tissue culture plastic (TCP). NEAT1 expression, quantified using qRT-PCR, revealed a significant increase on softer, brain-mimicking substrates compared to stiffer TCP. Overexpression of NEAT1 in brain-mimicking stiffness also showed higher expression levels of cancer progression markers, quantified by qRT-PCR. To further explore NEAT1’s role in oncogenesis due to its mechano-sensing capabilities, siRNA-mediated downregulation of NEAT1 in GBM cells cultured on soft PAA gels was performed to reduce NEAT1 levels to those comparable to TCP. The results showed reduced cancer aggressiveness, characterized by decreased expression of invasion, stemness, and epithelial-mesenchymal transition (EMT) markers, further supported by migration and invasion assays. These findings highlight NEAT1 as a mechanosensitive regulator in response to substrate stiffness, underscoring its role in tumor progression.

## Introduction

Cancer arises from a complex interplay of genetic factors that disrupt the gene networks responsible for maintaining cellular homeostasis. While much of cancer research has focused on mutations in protein-coding genes, emerging evidence highlights the significant role of non-coding RNAs (ncRNAs) in cancer biology^1^. Among these, long non-coding RNAs (lncRNAs)— transcripts longer than 200 base pairs—have gained prominence due to their diverse regulatory roles in cellular processes^2,3^. Unlike protein-coding genes, lncRNAs function as molecular scaffolds, guides, and decoys, influencing gene expression through interactions with DNA, RNA, and proteins^4,5^. In cancer, lncRNAs often exhibit dysregulated, tissue-specific expression and play key roles in processes such as cell cycle regulation, survival, immune response, metastasis, and stemness maintenance^2,3^. One of the most well-studied lncRNAs, nuclear paraspeckle assembly transcript 1 (NEAT1), has emerged as a pivotal oncogenic factor implicated in several solid tumors, including liver, prostate, gastric cancers, and gliomas^6–15^. However, the regulatory mechanisms governing NEAT1’s expression in cancer, particularly in response to the tumor microenvironment (TME), remain poorly understood.

The TME is a dynamic entity that significantly influences tumor progression and therapeutic resistance. One of its key characteristics is mechanical alteration, particularly the stiffening of the extracellular matrix (ECM) as tumors grow and remodel their surroundings^16^. This tissue stiffening, a hallmark of many solid tumors, regulates a wide range of cellular behaviors, including migration, proliferation, and gene expression. Stiffening induces changes in mechano-transduction pathways, such as those involving integrins, YAP/TAZ signaling, and cytoskeletal proteins, which convert mechanical stimuli into biochemical signals that promote cancer progression^17–19^. While considerable research has examined how substrate stiffness influences protein-coding gene expression, little is known about how mechanical cues affect the non-coding transcriptome, particularly lncRNAs.

This knowledge gap is critical because lncRNAs, unlike mRNAs, have unique regulatory mechanisms that may respond differently to environmental cues. Exploring how lncRNAs such as NEAT1 are influenced by substrate stiffness could provide crucial insights into how tumors adapt to mechanical changes in their environment and how this adaptation impacts disease progression. Given the diverse roles of lncRNAs in chromatin remodeling, transcriptional regulation, and RNA processing, their mechano-sensitivity could reveal novel regulatory networks that contribute to tumor aggressiveness, immune evasion, and metastatic potential. Targeting mechano-transduction pathways that influence NEAT1 expression could offer new therapeutic strategies for modulating the mechanical aspects of tumor biology to inhibit cancer progression.

Various different cancer types, including GBM have been shown to exhibit pronounced sensitivity to substrate stiffness, which plays a critical role in cancer progression^20–23^. A previous study from our group has also demonstrated that GBM cells are highly responsive to the mechanical properties of their environment, with softer, brain-like matrices promoting more aggressive tumor behavior and enhancing the invasive characteristics of relapsed GBM cells—a phenomenon not observed in conventional tissue culture plastic (TCP) models^24^. Additionally, the oncogenic potential of lncRNA NEAT1 has been extensively studied in GBM, with its involvement in pathways related to proliferation, epithelial-to-mesenchymal transition (EMT), and drug resistance^11,15,25–27^. However, whether the oncogenic nature of this lncRNA is influenced by its mechano-sensing abilities remains unexplored.

Our study addresses this gap by investigating the regulation of NEAT1 in glioblastoma (GBM) cells cultured on polyacrylamide (PAA) gels with brain-like stiffness (0.5kPa) versus the stiffer substrate of conventional tissue culture plastic (TCP). By mimicking the mechanical properties of the brain, we examined how substrate stiffness affects NEAT1 expression and functionality. qRT-PCR analysis revealed that NEAT1 expression was significantly elevated—up to 3-fold— on softer, brain-mimicking substrates compared to the stiffer TCP. This finding suggests that NEAT1 plays a role in mechano-sensing within different cancer micro-environments. Further experiments using siRNA-mediated NEAT1 downregulation in PAA gels with brain-like stiffness demonstrated a reduction in aggressive cancer characteristics, including decreased invasion, stemness, and epithelial-mesenchymal transition (EMT) markers. Functional assays, such as migration and invasion tests using Boyden’s chamber, validated these results. These findings highlight NEAT1’s contribution to GBM progression in response to substrate stiffness, positioning it as a potential target for mechano-sensitive therapeutic strategies in cancer treatment.

Our study not only provides novel insights into the mechano-regulation of lncRNAs but also identifies potential therapeutic approaches targeting NEAT1’s mechano-sensitivity to disrupt tumor progression. Understanding the role of the long non-coding transcriptome in response to the evolving mechanical properties of the TME is crucial for developing personalized and effective cancer therapies.

## Materials and method

### Correlation Analysis

The correlation analysis was carried out in bulk patient tumor RNAseq data deposited in The Cancer Genome Atlas (TCGA) using the “Correlation analysis” function in GEPIA 2^28^. The correlation of the lncRNAs NEAT1 was carried out with four pathways-Rho-GTPase signalling, HIPPO signalling, WNT B-Catenin and YAP-TAZ signalling. The signatures of the pathways were obtained from MSigDB^29^. The reported correlation coefficients were estimated using Spearman correlation method.

### PAA gel formation

Polyacrylamide (PAA) gel solutions with defined stiffness were prepared following the protocol outlined by Justin Tse et al.^30^. In summary, Acrylamide (Sigma-Aldrich) was combined with bisacrylamide (Sigma-Aldrich) in varying volumes. Milli-Q water was added to adjust the total volume. Polymerization was initiated by adding 1/100th of the total volume of 10% Ammonium persulfate (APS) (Sigma-Aldrich) and 1/1000th of the total volume of N,N,N’,N’-Tetramethylethylenediamine (TEMED) (Sigma-Aldrich).

### Cell Culture

U87-MG cells (an aggressive glioblastoma cell line, generously provided by Prof. Shilpee Dutt’s Lab at ACTREC, Navi Mumbai) were used in this study. They were maintained in DMEM (Gibco) supplemented with 10% FBS (HiMedia), 10% antibiotic-antimycotic (Anti-Anti, Gibco), and 1% L-Glutamine (Gibco). The cells were incubated in a humidified environment at 37°C with 5% CO2. Cells were seeded on plastic coverslips (22 mm^2^ squares) and on PAA gels prepared on 22 mm^2^ coverslips with designated stiffness, at a density of 100,000 cells. The cells were harvested after 48 hours of incubation, depending on the experiment.

### Treatments and transfections

For transfection and treatments, cells were plated in 12-well plates at densities of 4 × 104 cells per well, and allowed to incubate for 24 hours. For siRNA transfections, 15 nM siRNA was introduced into cells plated in 12-well plates. The transfected cells were maintained in Opti-MEM™, a reduced serum medium, for 4 hours, after which the medium was replaced with complete DMEM supplemented with FBS. For treatment of U87-MG cells with different inhibitor molecules, each inhibitor was administered at the following concentrations: Blebistatin: 10 μM, Y27: 50 μM, Latrunculin B: 3 μM, Nocodazole: 10 μM, and LPA: 20 μM. Post transfection and treatments as mentioned above, the cells were then incubated for 48 hours before harvesting them for subsequent experimental procedures.

### RNA Extraction and cDNA Synthesis

RNA was extracted from the harvested cells using TRIZOL® reagent (Ambion®), following the manufacturer’s step-by-step protocol. Total RNA isolation was performed according to the provided instructions. For each sample, cDNA was synthesized from 1 µg of RNA using the Biorad iScript™ cDNA synthesis kit, in line with the manufacturer’s guidelines.

### qRT-PCR

After cDNA preparation, real-time qPCR was performed on all samples, with technical duplicates for each. The reaction volume was 15 µl, including 1 µl of cDNA (diluted 1:2). The PCR conditions were as follows: initial denaturation at 95°C for 3 minutes, followed by 40 cycles of these steps: 95°C for 10 seconds, 60°C for 30 seconds, and 72°C for 30 seconds. The qRT-PCR primers used for various experiments are listed in Supplementary Table 1. Transcript quantification was carried out using iTaq™ Universal SYBR® Green Supermix (Biorad) on the QuantStudio™ 5 Real-Time PCR instrument (Thermo). The Ct values of each transcript were normalized to GAPDH. The fold change in transcript expression for comparative analysis was calculated using the 2-ΔΔCt method^31,32^. In brief, the fold change for each transcript was determined using the following formula:

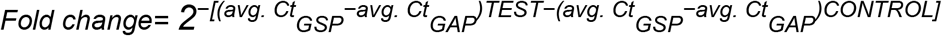

where avg. refers to average Ct values, GSP refers to gene-specific primers, GAP refers to GAPDH-specific primers, and TEST and CONTROL represent the respective experimental conditions.

### Migration and Invasion assay using Boyden’s chamber

Migration and invasion assays for the harvested cells from different substrates were performed using Boyden’s chambers. Briefly, cells were counted and seeded onto trans-well inserts at a density of 1.5 × 105 cells for both migration and invasion assays in FBS-free DMEM. The cells were allowed to migrate or invade for 24 hours at 37°C in a humidified incubator. The 24-well plates, on which the trans-well inserts were placed, contained complete media with FBS as a chemoattractant. For both the migration and invasion assays, the trans-wells had a permeable polycarbonate basement membrane. Additionally, for the invasion assay, the trans-wells were coated with a layer of Matrigel (1 mg/ml) to simulate the extracellular matrix (ECM). After the 24-hour incubation, non-migrating or non-invading cells on the top of the inserts were removed. The cells that had migrated or invaded through the membrane to the other side were fixed with methanol, stained with crystal violet dye, and imaged using an EVOS FL microscope with a 10X objective. The dye from the stained cells on the membrane was extracted with methanol, and the optical density at 590 nm was measured using a TECAN plate reader for quantification.

### Statistics

Statistical analysis was performed using GraphPad Prism 8.0 and Microsoft Excel to evaluate the significance across experimental replicates. Data are presented as mean ± S.D. or ± S.E., as noted in the figure legends, and derived from three independent biological replicates unless stated otherwise. A two-tailed unpaired Student’s t-test was employed for statistical comparisons, with a p-value < 0.05 considered significant. Statistical significance is represented as follows: * for P < 0.05, ** for P < 0.01, *** for P < 0.001, and **** for P < 0.0001. The biological replicates adhered to standard protocols in the field.

## Results

### Establishing the correlation between NEAT1 and mechano-sensing pathways from TCGA GBM patient samples

The long non-coding RNA NEAT1 has been extensively studied for its oncogenic potential and is implicated in various cancers, including breast, lung, colorectal, and glioblastoma (GBM). Its dysregulation is associated with critical tumor characteristics such as growth, invasion, metastasis, and resistance to treatment, positioning NEAT1 as a promising biomarker and therapeutic target. NEAT1 promotes cancer progression by regulating gene expression and cellular pathways related to stemness, proliferation, apoptosis, epithelial-mesenchymal transition (EMT), and DNA damage response, highlighting its significant role in cancer development and therapeutic resistance.

Despite extensive research on NEAT1’s oncogenic roles, a significant gap persists in understanding how substrate stiffness affects its expression. In the context of cancer, the evolving tumor microenvironment (TME) is a perfect example where oncogenic lncRNAs like NEAT1, if mechano-sensing in nature, can have a constant change in expression. Therefore, understanding this might be crucial to know how the mecchano-sensing lncRNAs orchestrate the regulatory mechanisms at different stages of the disease. Conventional tissue culture plastic (TCP) fails to capture these changes, making it essential to determine whether NEAT1 functions as a mechano-sensing lncRNA and whether its expression levels vary when cultured on substrates that mimic physiological tissue stiffness compared to TCP.

An earlier study from our lab demonstrated that NEAT1 exhibited the strongest positive correlation with hallmark cancer pathways, including p53, apoptosis, EMT, and proliferation, in GBM, based on TCGA patient data analysis. These findings underscore NEAT1’s clinical relevance across various cancer types, particularly in GBM. Furthermore, additional studies on GBM have highlighted NEAT1’s contribution to multiple facets of cancer biology that drive GBM progression. The mechano-sensitive nature of GBM cells has also been reported earlier through various studies. To investigate the potential role of NEAT1 in mechano-sensing pathways in glioblastoma (GBM) cells, we re-visited the TCGA database and analyzed data to examine correlations between NEAT1 expression and mechano-regulatory pathways in GBM patient samples. We identified a statistically significant moderate positive correlation between NEAT1 and several key mechano-sensing pathways, including Rho-GTPase, Wnt/β-Catenin, and Hippo signaling (Figure A(i–iii)). Notably, NEAT1 expression also correlated positively with YAP expression (Figure A(iv)), a central transcription factor in the Hippo pathway and a known oncogene implicated in cancer progression. YAP’s cellular localization is critical for mechano-sensing; it translocates into the nucleus on stiffer substrates but remains cytoplasmic on softer ones. Within the nucleus, YAP functions as a transcription factor for genes that mediate both mechanotransduction and cancer progression. However, in the context of GBM, the distinct impacts of YAP on these processes remain unclear. Thus, while the correlation in Figure A(iv) reveals an association between NEAT1 and YAP expression levels, it does not capture YAP’s localization within cells—a key aspect of its mechano-regulatory role. Investigating this cellular localization of YAP in GBM related to expression of NEAT1 could shed light on YAP’s mechanistic contributions to both lncRNA mediated mechano-sensing and oncogenesis, providing a compelling direction for future research.

In summary, NEAT1 plays a pivotal role in glioblastoma progression and is closely linked to key oncogenic pathways. However, further investigation into how substrate stiffness and the tumor microenvironment influence NEAT1 expression is essential for enhancing our understanding of its mechano-sensing capabilities and therapeutic potential.

### NEAT1 expression is increased in U87-MG GBM cells cultured on brain-like soft PAA gels when compared to conventional tissue culture plastic (TCP)

To evaluate whether NEAT1 functions as a mechano-sensing lncRNA and if its expression levels vary on substrates mimicking physiological tissue stiffness compared to tissue culture plastic (TCP), we created polyacrylamide (PAA) gels with a brain-like stiffness of 0.5 kPa. U87-MG GBM cells were grown on both the 0.5 kPa gel and TCP, and the expression of NEAT1 was quantified using qRT-PCR. We found that NEAT1 expression was elevated by up to 2.5-fold on the brain-mimetic substrate compared to TCP (Figure 2A), indicating that NEAT1 is mechano-sensing in nature.

To validate that the upregulation of NEAT1 on the 0.5 kPa gel was due solely to the mechanical environment and not due to difference in the chemical properties of the materials, we also cultured U87-MG cells on PAA gels with stiffness values of 5, 10, 40, and 100 kPa, in addition to the 0.5 kPa gel and TCP. The highest expression of NEAT1 was observed on the 0.5 kPa brain-like gel, exceeding a 3-fold increase compared to other stiffnesses tested (Figure 2B). Therefore, we conclude that NEAT1 lncRNA levels are sensitive to substrate stiffness, with the highest expression occurring when GBM cells are grown on substrates that mimic physiological brain-like stiffness.

### GBM cells with increased NEAT1 expression in brain-mimetic stiffness causes elevation in cancer-related mRNA levels when compared to TCP

In our previous study, we found that NEAT1 exhibited the highest positive correlation with GBM patient cancer samples across various cancer pathways (Figure 1B (i)). Our qRT-PCR studies revealed that NEAT1 expression levels in U87-MG GBM cells were upregulated by 3-fold when cultured on brain-mimetic 0.5 kPa stiffness gels compared to stiffer TCP substrates (Figure 2A (iv)). Therefore, we aimed to determine whether U87-MG cells grown on softer substrates displayed more aggressive cancer characteristics than those on stiffer TCP substrates, and whether this behavior was attributed to the mechano-sensing nature of NEAT1, with increased expression on the softer substrates.

**Figure 1.**
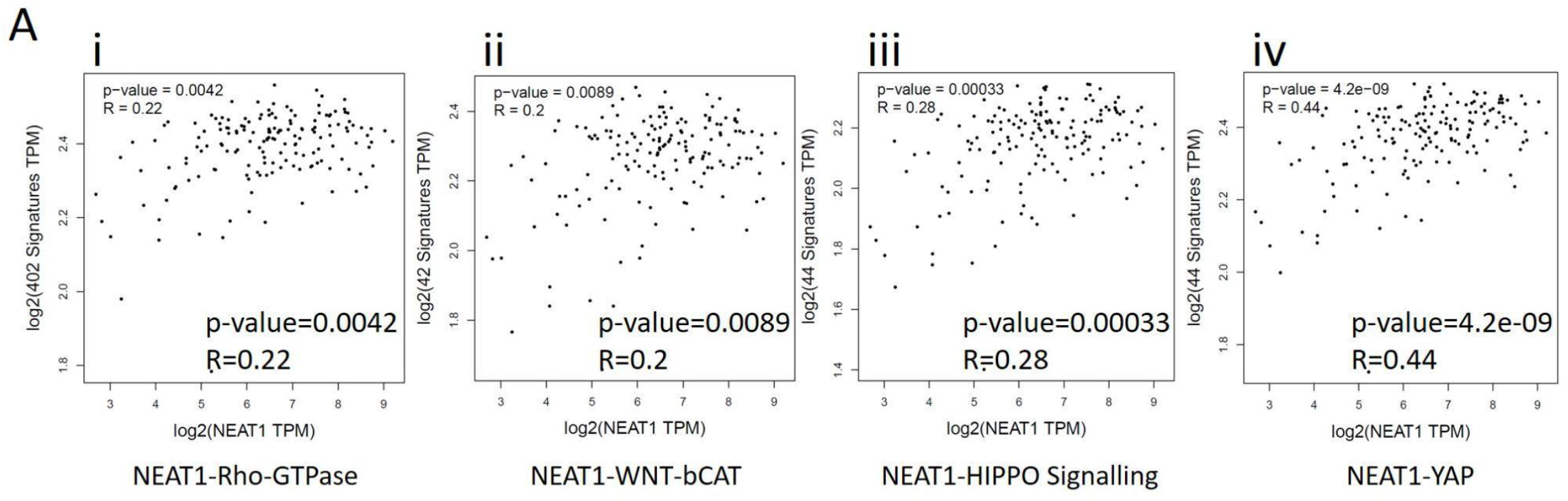
Establishing the correlation between NEAT1 and mechano-sensing pathways from TCGA GBM patient samples. A(i) Correlation of NEAT1 to Rho-GTPase pathway. (ii) Correlation of NEAT1 Wnt-Beta Catenin Pathway. (iii) Correlation of NEAT1 to HIPPO signalling pathway. (iv) Correlation of NEAT1 to YAP transcription factor involved in HIPPO signalling pathway.

**Figure 2.**
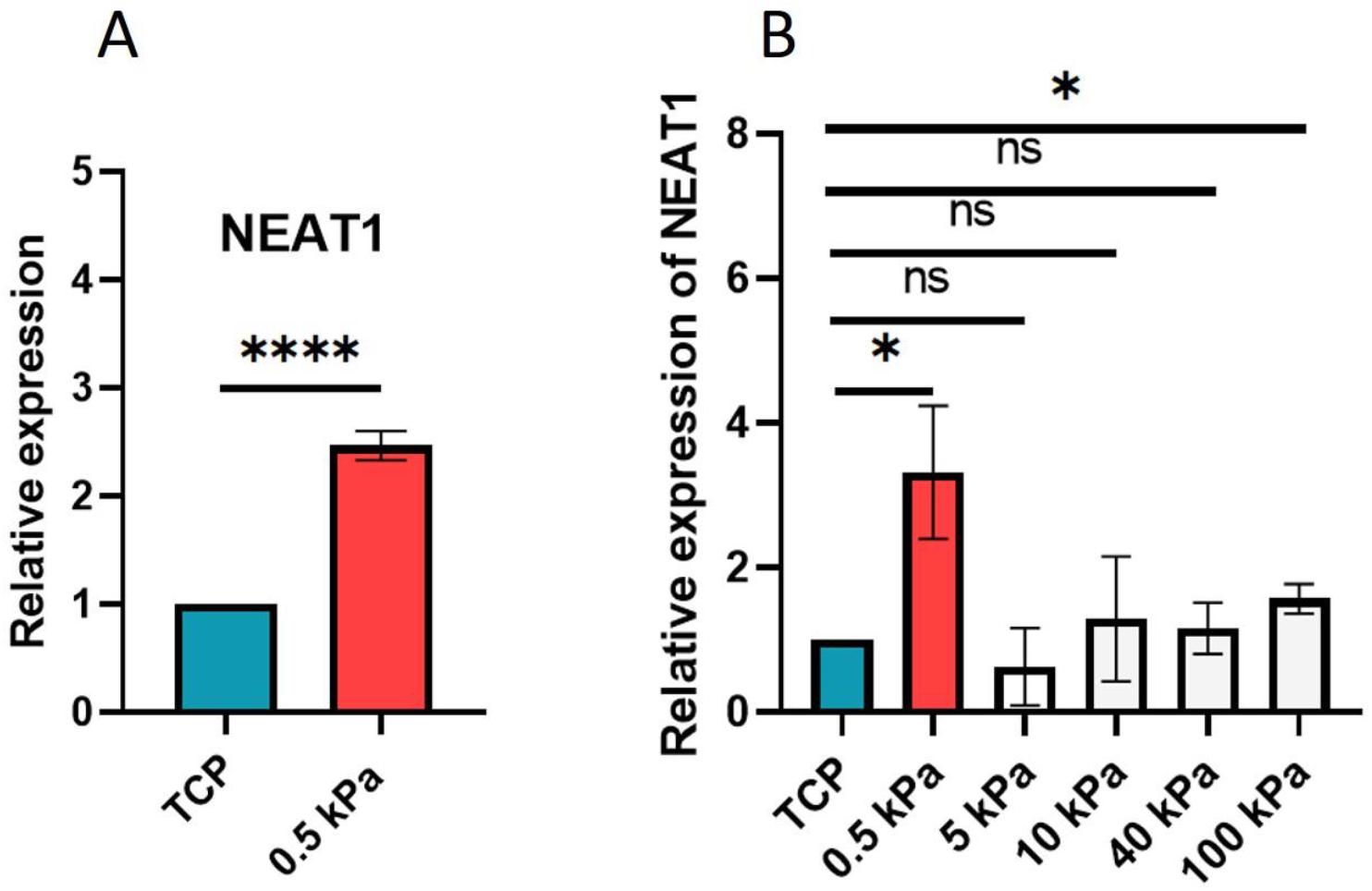
NEAT1 expression is increased in U87-MG GBM cells cultured on brain-like soft PAA gels when compared to conventional tissue culture plastic (TCP). (A) Relative expression levels of lncRNA NEAT1 in U87-MG cells grown on brain-mimetic 0.5kPa stiffness compared to TCP. (B) Relative expression levels of lncRNA NEAT1 in U87-MG cells grown on TCP and also on varying stiffness of PAA gels ranging from 0.5kPa brain-like stiffness to 100kPa. Error bars in (A and B) represent ±S.D. across three independent biological replicates. *P<0.05, **P<0.01, ***P<0.001, and ****P<0.0001.

To investigate this, we designed qRT-PCR experiments targeting cancer pathway genes associated with NEAT1, including stemness markers (OCT4, SOX2, NANOG, CD133), invasion markers (MMP2, MMP9, Fibronectin), glucose transporters (GLUT1, GLUT3), and EMT-related genes (Snail, Slug, Vimentin, N-Cadherin, Beta-catenin) (Supplementary Table 1). We first measured the NEAT1 expression levels in U87-MG cells cultured on soft 0.5 kPa gels and TCP. To validate that the resultant changes in the cancer markers is solely due to the NEAT1 expression variation, we used siRNA mediated NEAT1 downregulation for cells grown on the 0.5 kPa brain-like stiffness. Given NEAT1’s oncogenic nature, we aimed to ascertain whether the variations in the cancer-related behavior of U87-MG cells between 0.5 kPa and TCP were influenced only by substrate stiffness. Therefore, we downregulated NEAT1 in the GBM cells grown on 0.5 kPa gels using siRNAs, reducing its expression to levels comparable to those in cells grown on TCP (Figure 3A).

**Figure 3.**
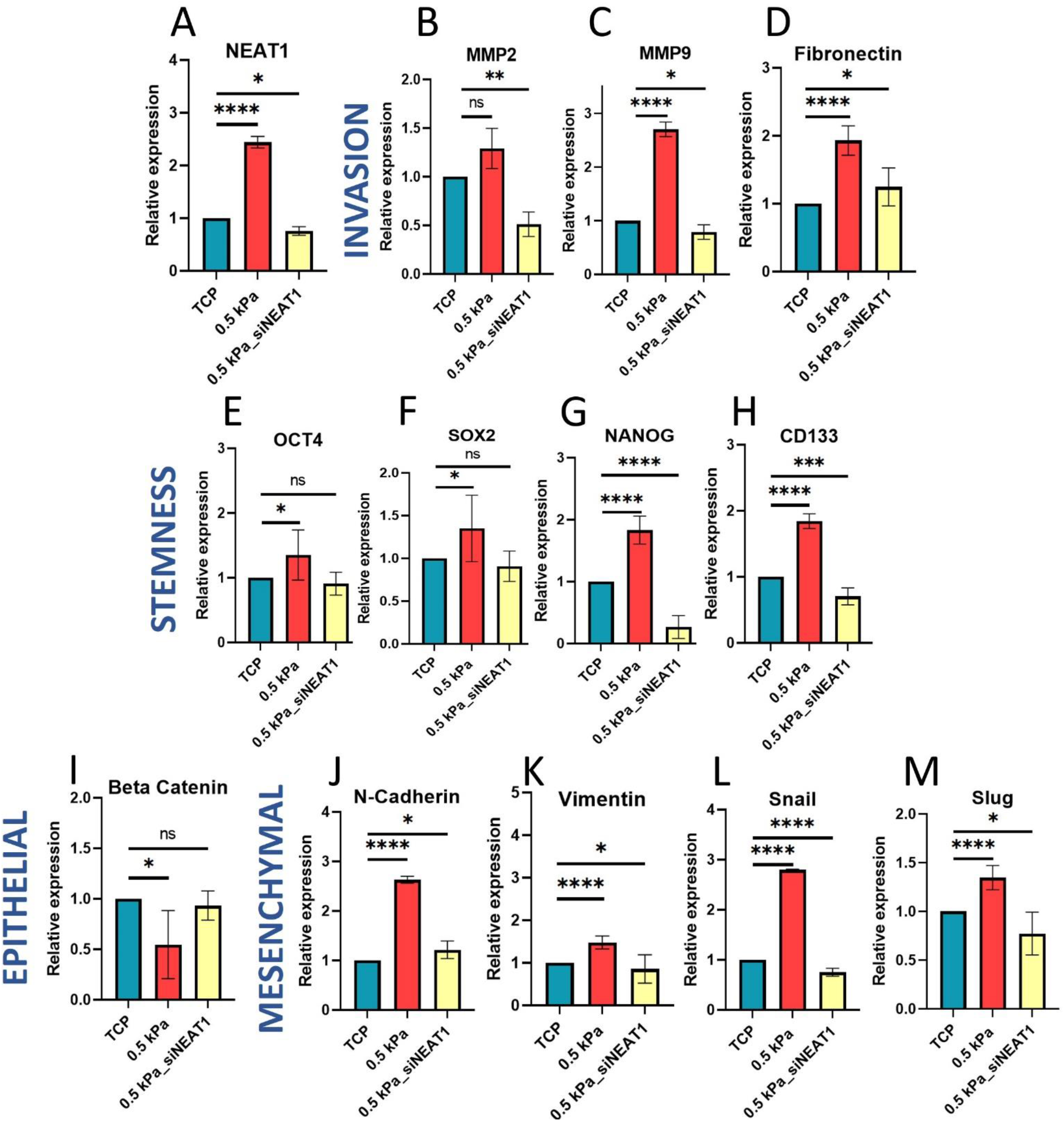
GBM cells with increased NEAT1 expression in brain-mimetic stiffness causes elevation in cancer-related mRNA levels when compared to TCP. (A) Relative expression levels of NEAT1 in U87-MG cells grown on brain-like (0.5kPa) stiffness, TCP and, and NEAT1-downregulated cells (via siRNA) on brain-mimetic stiffness. (B-D) Relative expression of invasion markers (Fibronectin, MMP2 and MMP9) in cells grown on brain-like (0.5kPa) stiffness, TCP and, and NEAT1-downregulated cells (via siRNA) on brain-mimetic stiffness. (E-H) Relative expression of stemness markers (OCT4, SOX2, NANOG and CD133) in U87-MG cells grown on brain-like (0.5kPa) stiffness, TCP and, and NEAT1-downregulated cells (via siRNA) on brain-mimetic stiffness. (I) Relative expression of epithelial marker (Beta Catenin) in U87-MG cells grown on brain-like (0.5kPa) stiffness, TCP and, and NEAT1-downregulated cells (via siRNA) on brain-mimetic stiffness. (J-M) Relative expression of mesenchymal markers (N-Cadherin, Vimentin, Snail and Slug) in U87-MG cells grown on brain-like (0.5kPa) stiffness, TCP and, and NEAT1-downregulated cells (via siRNA) on brain-mimetic stiffness. Error bars in (A-O) represent ±S.D. across three independent biological replicates. *P<0.05, **P<0.01, ***P<0.001, and ****P<0.0001.

Subsequently, we compared the relative mRNA expression of various cancer markers among cells cultured on soft gels, TCP, and NEAT1-downregulated cells on the soft substrate. We observed that invasion markers (MMP2, MMP9, Fibronectin) were markedly overexpressed in cells grown on soft substrates (Figure 3B-D). For instance, the invasion marker MMP2 did not show significant upregulation on the soft substrate compared to TCP (Figure 3B), but upon NEAT1 downregulation to levels equivalent to those in TCP, MMP2 expression was reduced by more than 50% on the soft gel. MMP9 also exhibited significant upregulation by more than 2.5-fold on soft substrates, and its expression decreased to levels comparable to TCP in NEAT1-downregulated cells (Figure 3C). Fibronectin was upregulated by approximately 2-fold in cells on 0.5 kPa gels compared to TCP, returning to almost TCP-like levels following NEAT1 downregulation (Figure 3D). This strong correlation between NEAT1 and invasive potential highlights the role of substrate stiffness in influencing GBM cell behavior.

Further, the stemness markers (OCT4, SOX2, NANOG, CD133) were also significantly elevated in cells grown on 0.5 kPa gels compared to TCP, with a more pronounced effect observed on NANOG and CD133 (Figure 3E-H). Upon NEAT1 downregulation on the softer substrate, the elevated expression of these markers was largely reversed (Figure 3E-H). For instance, OCT4 and SOX2 were upregulated by approximately 1.5-fold each on the soft gels compared to TCP, but both markers displayed mRNA levels equivalent to those on TCP following NEAT1 downregulation. NANOG exhibited a 2-fold upregulation on the soft substrate compared to TCP but, interestingly, showed expression levels 80% lower than those on TCP in NEAT1-downregulated cells (Figure 3G). The expression of CD133, which was upregulated by 2-fold on the softer substrates, reverted to levels similar to TCP after NEAT1 downregulation (Figure 3H). These results indicate that NEAT1 may significantly promote stemness in GBM cells cultured on softer substrates, a behavior that is likely more accurately captured in this context than in stiffer TCP environments.

Additionally, we observed that the epithelial marker Beta-Catenin was downregulated (Figure 3C), while mesenchymal markers (Vimentin, N-Cadherin, Snail, Slug) were upregulated in cells cultured on soft substrates compared to TCP (Figure 3J-M). Notably, Beta-Catenin expression in NEAT1-downregulated cells on the soft gel increased from 0.5-fold to 1-fold, comparable to levels observed in TCP cultures (Figure 3I). For mesenchymal markers, the greatest variation was observed in N-Cadherin expression, which declined from nearly 3-fold to levels equivalent to TCP (Figure 3J). Vimentin levels showed significant variability across the soft gel, TCP, and NEAT1-downregulated cells, but the differences among the three were minimal (Figure 3K). An effect similar to N-Cadherin was observed for Snail (Figure 3L). Slug was mildly upregulated on the soft gel, with expression increasing up to 1.5-fold, but after NEAT1 downregulation, it decreased by 20% compared to TCP levels (Figure 3M). These findings suggest that NEAT1 plays a crucial role in modulating epithelial-mesenchymal transition (EMT) in GBM, with downregulation in cells on softer substrates effectively reflecting dynamic changes in EMT-related markers, providing valuable insights into cancer progression and metastasis.

Overall, our results highlight that NEAT1 significantly influences various aspects of GBM biology, including stemness, invasion, and EMT, with its effects being more pronounced in cells cultured on softer, brain-like substrates compared to stiffer TCP. This underscores the importance of investigating lncRNAs like NEAT1 in microenvironments that mimic physiological tissue stiffness to gain deeper insights into tumor aggressiveness and identify potential therapeutic targets.

### Higher NEAT1 expression in GBM cells on 0.5 kPa brain-like stiffness causes increased migration and invasion when compared to cells grown on TCP

Our studies show that the mRNA expression of various cancer markers especially of invasion and EMT have a correlation with NEAT1 expression. Therefore, we conducted functional assays to analyze whether those effects could be captured at a phenotypic level. To assess the migration and invasion potential of GBM cells under different substrate stiffnesses, we employed the Boyden’s chamber assay. Cells from both soft and stiff substrates were seeded on the transwell inserts, with the invasion inserts coated in Matrigel to simulate extracellular matrix barriers, which are critical for invasion as opposed to migration (Figure 4A,D). After 24 hours, migrated or invaded cells were stained with crystal violet, visualized, and quantified by measuring optical density at 590 nm. Cells from the softer brain-like 0.5 kPa PAA gels demonstrated a 2-fold increase in migration potential compared to those from the stiffer TCP (Figure 4B,C). However, the migration capacity of NEAT1 downregulated cells cultured on soft substrates was reduced. For invasion, measured through the ability to penetrate a Matrigel layer in the Boyden’s chamber, cells on the softer substrates showed a significant 0.5-fold increase in invasion compared to those on stiff TCP (Figure 4E,F). Furthermore, NEAT1 downregulation in cells from the soft substrates resulted in invasion levels comparable to those on stiffer substrates (Figure 4E,F).

**Figure 4.**
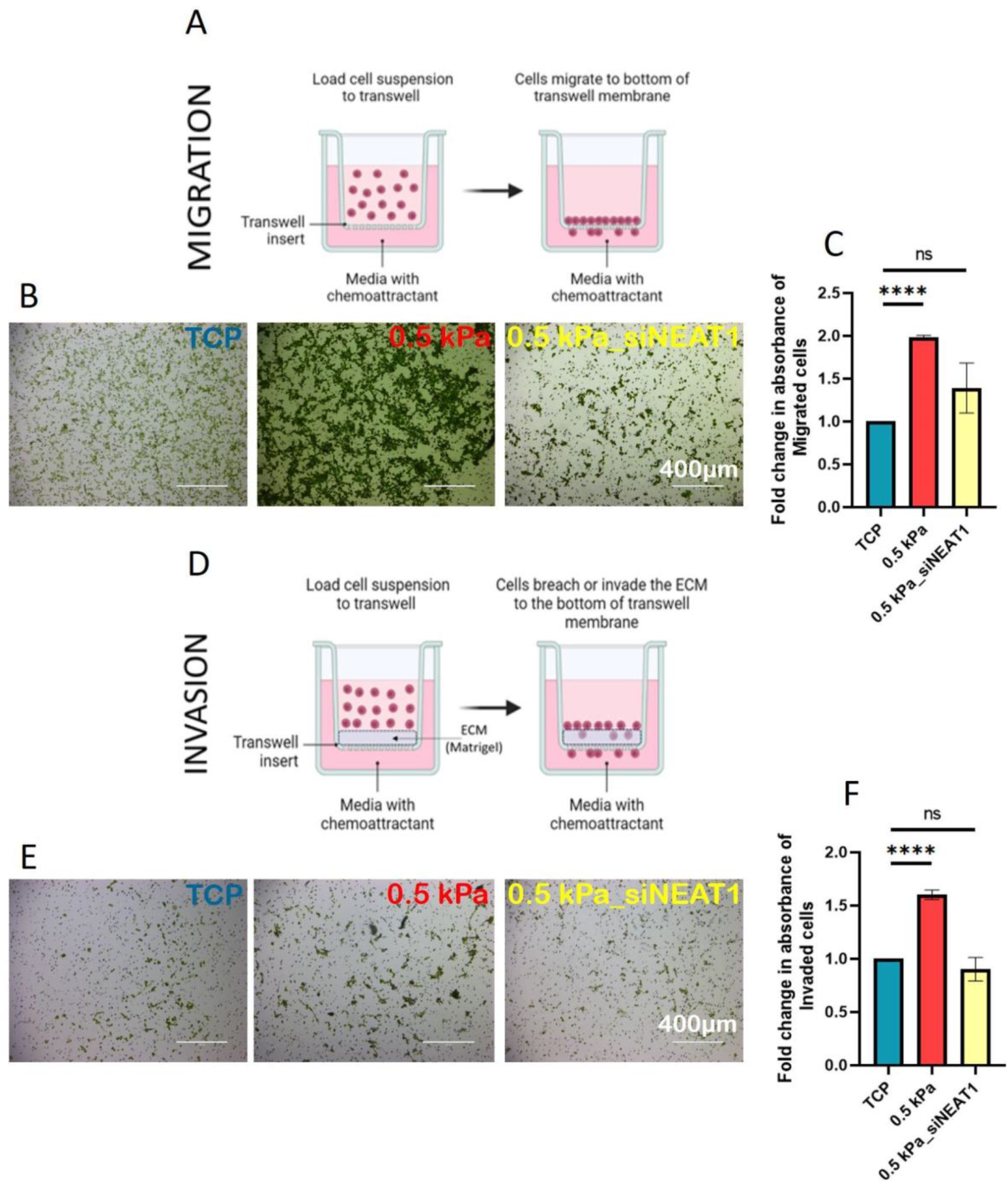
Higher NEAT1 expression in GBM cells on 0.5 kPa brain-like stiffness causes increased migration and invasion when compared to cells grown on TCP. (A) Schematic representation to describe the process of migration through the Boyden’s chamber. (B) Representation of cells from brain-like (0.5kPa) stiffness, TCP and, and NEAT1-downregulated GBM cells on brain-like stiffness, which migrated through the trans-well chambers (without Matrigel). Scale bar corresponds to 400µm. (C) Fold change in the absorbance of the dye at 560 nm, extracted from the cells that migrated the trans-well chamber denoting the comparison in the motile properties of the cells from brain-like (0.5kPa) stiffness, TCP and, and NEAT1-downregulated GBM cells on brain-like stiffness. (D) Schematic representation to describe the process of invasion through the Boyden’s chamber. (B) Representation of cells from brain-like (0.5kPa) stiffness, TCP and, and NEAT1-downregulated GBM cells on brain-like stiffness, which invaded through the Matrigel coated trans-well chambers. Scale bar corresponds to 400µm. (C) Fold change in the absorbance of the dye at 560 nm, extracted from the cells that invaded the trans-well chamber denoting the comparison in the invasive properties of the cells from brain-like (0.5kPa) stiffness, TCP and, and NEAT1-downregulated GBM cells on brain-like stiffness. Error bars represent ±S.E.; *P<0.05, **P<0.01, ***P<0.001, and ****P<0.0001.

These findings underscore that elevated NEAT1 levels in cells cultured on softer substrates significantly influence the aggressive characteristics of GBM cells when compared to those on stiffer substrates. The increased variability in proliferation rates among cells on softer substrates, combined with the substantial impact of NEAT1 downregulation on migration, and invasion, highlights NEAT1’s critical role in modulating the aggressive behavior of GBM cells within a mechanically responsive environment.

### NEAT1 expression is upregulated by actin-myosin destabilization and downregulated by microtubule disruption

From our observation, it is evident that NEAT1 functions as a mechano-sensing lncRNA in GBM, with its elevated expression on soft, brain-mimetic stiffness contributing to a more aggressive GBM phenotype, a pattern not observed on TCP. To investigate which cellular mechano-sensing pathways are involved in the upregulation of NEAT1 on different substrates, we treated cells on TCP with various pathway inhibitors (Figure 5A): actin inhibitor— Latrunculin B, myosin inhibitor—Blebbistatin, ROCK inhibitor—Y27632 (Y27), microtubule inhibitor—Nocodazole, and GPCR inhibitor—Lysophosphatidic acid (LPA). Cells grown on stiff substrates typically exhibit increased focal adhesions, enhanced cytoskeletal tension, and greater activation of mechanosensing pathways, promoting a more spread-out morphology and higher proliferation rates. In contrast, cells on soft substrates tend to have reduced focal adhesion, lower cytoskeletal tension, and a more rounded morphology, which leads to decreased mechanosensing activity and slower proliferation. In a stiffer TCP, Latrunculin B and Blebbistatin, which inhibit actin and myosin polymerization, respectively, both resulted in a 2-fold increase in NEAT1 levels, although Blebbistatin’s effect was statistically significant and more consistent. The ROCK inhibitor Y27 also led to an increase in NEAT1 expression, albeit inconsistently. The GPCR inhibitor LPA had no effect on NEAT1 expression. On the contrary, the microtubule inhibitor Nocodazole had an opposite effect, reducing NEAT1 expression by more than 70%. Next, we analyzed the effect of these inhibitors on GBM cells grown on a soft 0.5 kPa gel, where NEAT1 expression is inherently elevated compared to TCP. On this soft substrate, Latrunculin B and Blebbistatin had no effect on NEAT1 levels (Figure 5B). Y27 and LPA showed a non-significant downregulation of NEAT1. However, Nocodazole treatment again significantly downregulated NEAT1, by up to 90%. Intrigued by this, we further investigated the combined effects of Nocodazole and Blebbistatin on the 0.5 kPa gel. We found that, despite Nocodazole downregulating NEAT1, Blebbistatin treatment led to an upregulation (Figure 5C). Similarly, when NEAT1 was downregulated using siRNA on the soft gel, Blebbistatin treatment resulted in NEAT1 upregulation once again (Figure 5C).

**Figure 5.**
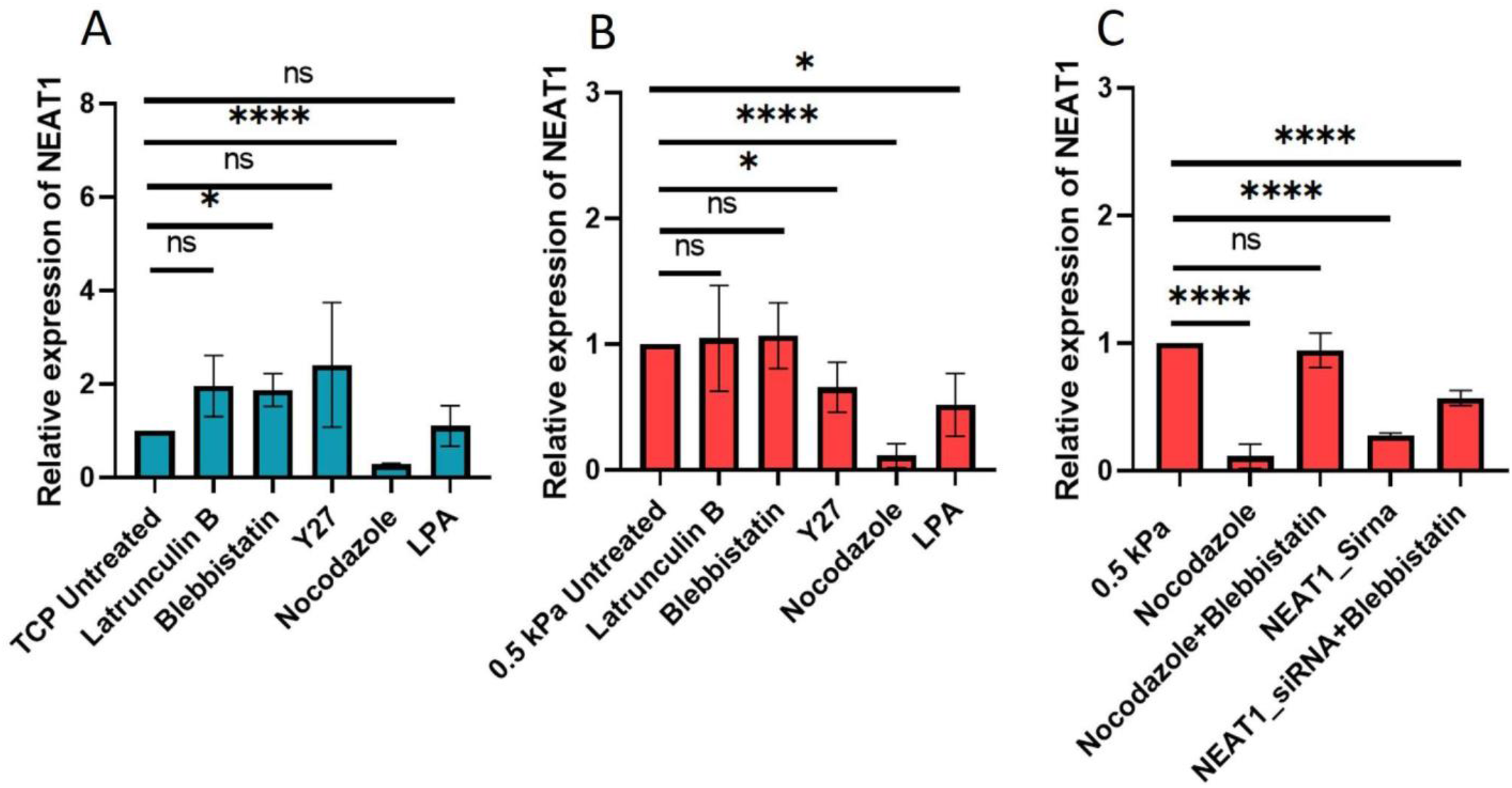
NEAT1 expression is upregulated by actin-myosin destabilization and downregulated by microtubule disruption. (A) Relative expression of NEAT1 levels in U87-MG cells grown on TCP (untreated) compared to cells treated with Latrunculin B, Blebbistatin, Y27, Nocodazole and LPA. (B) Relative expression of NEAT1 levels in U87-MG cells grown on brain-like 0.5kPa stiffness (untreated) compared to cells treated with Latrunculin B, Blebbistatin, Y27, Nocodazole and LPA. (C) Relative expression of NEAT1 levels in U87-MG cells grown on brain-like 0.5kPa stiffness (untreated) compared to cells grown on brain-like 0.5kPa stiffness and treated with Nocodazole, Blebbistatin, Nocodazole followed by Blebbistatin, and NEAT1-downregulated cells (via siRNA) followed by Blebbistatin treatment. Error bars represent ±S.E.; *P<0.05, **P<0.01, ***P<0.001, and ****P<0.0001.

In conclusion, NEAT1 upregulation is linked to actin and myosin inactivation, while microtubule disengagement leads to its downregulation. Therefore, these mechano-transduction molecules play a crucial role in modulating NEAT1 levels on soft and stiff substrates.

## Discussion

LncRNAs have emerged as critical regulators in cancer biology, influencing gene expression, tumor progression, metastasis, and therapy resistance^2,3^. NEAT1, a prominent oncogenic lncRNA, plays a significant role in various solid tumors, including GBM, by participating in pathways that drive cellular proliferation, invasion, EMT, and chemoresistance^6–15^. However, the differential response of lncRNAs like NEAT1 to the mechanical properties of the ECM, particularly stiffness, remains underexplored^33,34^. While the effects of substrate stiffness on gene and protein dysregulation have been thoroughly studied^16–19^, the impact on the long non-coding genome under different physiological or pathological conditions is largely unknown. Some research has examined mechano-sensing lncRNAs in endothelial cells, identifying lncRNAs such as LASSIE, HOTAIR, ANRIL, and MEG3^35–38^ as responsive to mechanical changes. In cancer, the evolving TME and the associated changes in tissue stiffness over time raise important questions about their effects on lncRNA expression, which are yet to be addressed.

Two studies, by Todorovski et al. (2020) and Xu et al. (2021), report mechano-sensing lncRNAs in specific cancer types^33,34^. The former study suggests that NEAT1-associated paraspeckles act as mechano-sensors in breast cancer and osteosarcoma, forming more readily on softer substrates, indicating their role in mechano-sensitivity and mechano-memory in cancer mechanobiology. Although it does not address changes in NEAT1 lncRNA expression. The latter study demonstrates that NEAT1 is a mechanosensitive lncRNA regulating proliferation and EMT in liver cancer cells via the NEAT1/WNT/β-catenin pathway in response to increased ECM stiffness. From these studies, it is evident that cancer originating from different tissue types might have different effect on lncRNA expressions on varying substrate stiffnesses. Additionally, in a non-cancerous context, Liu et al. (2022) found that lncRNA NEAT1 and its paraspeckle structures are involved in osteoblast mechano-transduction, impacting bone formation and strength^39^. Despite these findings, there are currently no studies specifically investigating the effects of substrate stiffness on lncRNA expression and cancer progression in any other cancer types including GBM which is reported to be highly sensitive to substrate stiffness. Exploring these differences is vital for developing accurate models that replicate the TME and improve therapeutic strategies.

In an earlier study, we had conducted a correlation analysis using TCGA data to establish the clinical relevance of NEAT1 across various cancer types^40^. Our findings revealed significant positive correlations between NEAT1 expression and critical cancer pathways, including p53 signaling, apoptosis, epithelial-mesenchymal transition (EMT), and proliferation. The correlation was especially pronounced in GBM, indicating NEAT1’s central role in cancer progression. Consequently, we aimed to explore how mechanical cues, particularly matrix stiffness, affect NEAT1 expression and its contribution to cancer progression in GBM. To explore NEAT1’s role in mechano-sensing pathways, we revisited TCGA data to analyze correlations between NEAT1 expression levels and mechano-regulatory pathways in GBM patient samples^41–43^. We found a statistically significant positive correlation between NEAT1 and YAP, a transcription factor involved in the Hippo signaling pathway crucial for mechano-regulation. Additionally, NEAT1 exhibited moderate positive correlations with other mechano-regulatory pathways, including Wnt/β-Catenin and Rho-GTPase pathways.

To further investigate the impact of stiffness on NEAT1 expression, we cultured U87-MG glioblastoma (GBM) cells on polyacrylamide (PAA) gels with varying stiffness, mimicking the mechanical properties of normal brain tissue (soft) and tissue culture plastic (TCP). Quantitative real-time PCR (qRT-PCR) analysis revealed that NEAT1 expression was significantly upregulated by up to 2.5-fold in GBM cells cultured on the soft 0.5 kPa brain-mimetic matrix compared to those on TCP. To validate that this upregulation was due solely to the mechanical environment and not to differences in the chemical properties of the materials, we also analyzed NEAT1 expression in U87-MG cells cultured on PAA gels with stiffness values of 5, 10, 40, and 100 kPa. The highest NEAT1 expression was observed on the 0.5 kPa brain-like gel, exceeding a three-fold increase compared to other stiffnesses tested. These findings suggest that NEAT1 functions as a mechano-sensing lncRNA in GBM, with its expression levels being sensitive to substrate stiffness and maximum at its physiological stiffness, and, thus, might be contributing to enhanced cancer progression in brain-like mechanical environments.

To analyse the effect of elevated NEAT1 expression on the brain-like stiffness, through siRNA-mediated downregulation of NEAT1 in GBM cells grown on soft matrices, we reduced its expression to levels comparable to those on TCP. This downregulation resulted in significant changes in the expression of cancer-related markers. For example, the mRNA levels of stemness markers, including OCT4, SOX2, and NANOG, were markedly elevated in cells grown on soft 0.5kPa gels. Upon NEAT1 downregulation, these markers exhibited significantly reduced expression, indicating NEAT1’s role in maintaining cancer stemness in response to a soft matrix. The high stemness marker expression on soft matrices correlated with high NEAT1 expression aligns with previous studies linking cancer stemness to increased NEAT1 levels, drug resistance and tumor recurrence. Similarly, markers associated with invasion, such as MMP2 and Fibronectin, were significantly upregulated in GBM cells cultured on soft matrices. Their expression decreased upon NEAT1 downregulation, suggesting that NEAT1 promotes invasive properties in soft environments. Additionally, EMT markers, including Vimentin and N-Cadherin, followed a similar trend, with their expression levels being higher on soft matrices and reduced upon NEAT1 knockdown. The downregulation of NEAT1 also resulted in decreased expression of Snail, a master regulator of EMT, further implicating mechano-sensing role of NEAT1 in EMT-associated cancer progression in softer brain-like microenvironments.

To validate the observations regarding the mRNA expression of various cancer markers and their correlation with NEAT1 expression, we conducted functional assays to analyze migration and invasion in GBM cells cultured on soft 0.5kPa stiffness versus stiff TCP. To assess the migration and invasion potential of GBM cells under different mechanical conditions, we employed the Boyden’s chamber assay. Cells from the softer PAA gels demonstrated a 2-fold increase in migration potential compared to those from the stiffer TCP. However, the migration capacity of NEAT1 downregulated cells cultured on soft substrates was reduced. For invasion, measured through the ability of cells to penetrate a Matrigel layer in the Boyden’s chamber, cells on the softer substrates showed a significant 0.5-fold increase in invasion compared to those on stiff TCP. Furthermore, NEAT1 downregulation in cells from the soft substrates resulted in invasion levels comparable to those on stiffer substrates. These findings underscore that elevated NEAT1 levels in cells cultured on softer substrates significantly influence the aggressive characteristics of GBM cells when compared to those on stiffer substrates. The impact of NEAT1 downregulation on migration and invasion, highlights NEAT1’s critical role in modulating the aggressive behavior of GBM cells within a mechanically responsive environment.

Our experiments demonstrate that NEAT1 functions as a mechano-sensing lncRNA in GBM, with elevated expression on soft, brain-mimetic substrates correlating with a more aggressive GBM phenotype, unlike the expression pattern observed on stiff tissue culture plastic (TCP). To explore the mechano-sensing pathways involved in NEAT1 upregulation, we treated GBM cells on TCP with various pathway inhibitors, including Latrunculin B (actin inhibitor), Blebbistatin (myosin inhibitor), Y27632 (ROCK inhibitor), Nocodazole (microtubule inhibitor), and Lysophosphatidic acid (GPCR inhibitor)^44–48^. Both Latrunculin B and Blebbistatin resulted in a two-fold increase in NEAT1 levels, with Blebbistatin showing a statistically significant and consistent effect. The ROCK inhibitor Y27632 also increased NEAT1 expression, albeit inconsistently, while LPA had no effect. Conversely, Nocodazole reduced NEAT1 expression by over 70%. When analyzing cells on a soft 0.5 kPa gel, where NEAT1 is inherently elevated, Latrunculin B and Blebbistatin showed no impact on NEAT1 levels. This outcome was anticipated, as actin and myosin activity is already low on a soft matrix, resulting in minimal cell stretching and inherently high NEAT1 expression levels, likely approaching saturation. Consequently, additional inactivation of actin or myosin had no observable effect. Nocodazole treatment significantly downregulated NEAT1 by up to 90%. Notably, in combined treatments, Nocodazole downregulated NEAT1, but Blebbistatin treatment led to its upregulation. Similarly, downregulation of NEAT1 using siRNA on the soft gel also resulted in increased NEAT1 levels upon Blebbistatin treatment. These findings suggest that NEAT1 expression is intricately linked to the mechanical properties of the tumor microenvironment. The inactivation of actin and myosin pathways appears to enhance NEAT1 expression on TCP, while disengagement from microtubules reduces it both on TCP and soft brain-like stiffness. This relationship underscores the critical role of cytoskeletal dynamics in regulating NEAT1 levels, highlighting potential therapeutic targets for modulating GBM aggressiveness. By understanding the mechanistic pathways influencing NEAT1 expression, future strategies could aim to manipulate these pathways to improve treatment outcomes in GBM patients.

In conclusion, our study demonstrates that NEAT1 is a mechanosensitive lncRNA that mediates cancer progression in response to mechanical cues, particularly matrix stiffness. The upregulation of NEAT1 in stiffer environments enhances cancer stemness, invasion, EMT, and proliferation, implicating the importance of using models that accurately replicate the mechanical properties of tumors when studying lncRNA function. These findings highlight NEAT1’s potential as a therapeutic target, especially in aggressive cancers like GBM, where matrix stiffness significantly influences tumor behavior. Understanding how lncRNAs like NEAT1 are regulated by mechanical forces could pave the way for more effective cancer treatment strategies tailored to the mechanical properties of the TME. This opens avenues for future research on elucidating the specific mechano-regulatory pathways governing NEAT1 expression and their broader implications in developing targeted cancer therapies.

## Acknowledgements

We deeply acknowledge the laboratories of Prof. Shilpee Dutt (ACTREC Navi Mumbai) who kindly helped us by providing the desired cell line U87-MG for this study (as mentioned in the Methods section).

