## Supplementary Information for "Mechano-sensing role of oncogenic lncRNA NEAT1 on soft vs. stiff substrates and its implications for glioblastoma progression"

### Supplementary Table 1. List of qPCR Primers

| PRIMER NAME | SEQUENCE (5`-3`) |
|---|---|
| Vimentin FP | AGGCAAAGCAGGAGTCCACTGA |
| Vimentin RP | ATCTGGCGTTCCAGGGACTCAT |
| N-Cadherin FP | GCGTCTGTAGAGGCTTCTGG |
| N-Cadherin RP | GCCACTTGCCACTTTTCCTG |
| E-Cadherin FP | GGTTTTCTACAGCATCACCG |
| E-Cadherin RP | GCTTCCCCATTTGATGACAC |
| Beta Catenin FP | CACAAGCAGAGTGCTGAAGGTG |
| Beta Catenin RP | GATTCCTGAGAGTCCAAAGACAG |
| MALAT1 FP | GGGTGTTTACGTAGACCAGAACC |
| MALAT1 RP | CTTCCAAAAGCCTTCTGCCTTAG |
| NEAT1 FP | GCTGGACCTTTCATGTAACGGG |
| NEAT1 RP | TGAACTCTGCCGGTACAGGGAA |
| MMP2 FP | AGCGAGTGGATGCCGCCTTTAA |
| MMP2 RP | CATTCCAGGCATCTGCGATGAG |

| | |
|---|---|
| MMP9 FP | GCCACTACTGTGCCTTTGAGTC |
| MMP9 RP | CCCTCAGAGAATCGCCAGTACT |
| Fibronectin FP | ACAACACCGAGGTGACTGAGAC |
| Fibronectin RP | GGACACAACGATGCTTCCTGAG |
| CD133 FP | CACTACCAAGGACAAGGCGTTC |
| CD133 RP | CAACGCCTCTTTGGTCTCCTTG |
| Nanog FP | CTCCAACATCCTGAACCTCAGC |
| Nanog RP | CGTCACACCATTGCTATTCTTCG |
| Nestin FP | TCAAGATGTCCCTCAGCCTGGA |
| Nestin RP | AAGCTGAGGGAAGTCTTGGAGC |
| Glut1 FP | TTGCAGGCTTCTCCAACTGGAC |
| Glut1 RP | CAGAACCAGGAGCACAGTGAAG |
| Glut3 FP | CGTTGTTGGAATTCTGGTGGC |
| Glut3 RP | CTTAGCATTCTCCTCTTCTTTT |
| SOX2 FP | GCTACAGCATGATGCAGGACCA |
| SOX2 RP | TCTGCGAGCTGGTCATGGAGTT |
| OCT4 FP | CCTGAAGCAGAAGAGGATCACC |

| | |
|---|---|
| OCT4 RP | AAAGCGGCAGATGGTCGTTTGG |
| SLUG FP | ATCTGCGGCAAGGCGTTTTCCA |
| SLUG RP | GAGCCCTCAGATTTGACCTGTC |
| Snail FP | TGCCCTCAAGATGCACATCCGA |
| Snail RP | GGGACAGGAGAAGGGCTTCTC |
